# Nuclear arms races: experimental evolution for mating success in the mushroom-forming fungus *Schizophyllum commune*

**DOI:** 10.1101/479147

**Authors:** Bart P. S. Nieuwenhuis, Duur K. Aanen

## Abstract

When many gametes compete to fertilize a limited number of compatible gametes, sexual selection will favour traits that increase competitive success during mating. In animals and plants, sperm and pollen competition have yielded many interesting adaptations for improved mating success. In fungi, similar processes have not been shown directly yet. We test the hypothesis that sexual selection can increase competitive fitness during mating, using experimental evolution in the mushroom-forming fungus *Schizophyllum commune* (Basidiomycota). Mating in mushroom fungi occurs by donation of nuclei to a mycelium. These fertilizing ‘male’ nuclei migrate through the receiving ‘female’ mycelium. In our setup, an evolving population of nuclei was serially mated with a non-evolving female mycelium for 20 sexual generations. From the twelve tested evolved lines, four had increased and one had decreased fitness relative to an unevolved competitor. Even though only two of those five remained significant after correcting for multiple comparisons, for all five lines we found a correlation between the efficiency with which the female mycelium is accessed and fitness, providing additional circumstantial evidence for fitness change in those five lines. In two lines, fitness change was also accompanied by increased spore production. The one line with net reduced competitive fitness had increased spore production, but reduced fertilisation efficiency. We did not find trade-offs between male reproductive success and other fitness components. We compare these findings with examples of sperm and pollen competition and show that many similarities between these systems and nuclear competition in mushrooms exist.

## Introduction

Sexual selection is the component of natural selection associated with variation in mating success [1,2]. Sexual selection favours investment in traits that improve the likelihood of fertilization given limited access to opposite-sex gametes due to competition with members of the same sex [2] and is generally considered to be a significant component of natural selection [3]. Competition for mating can occur before mating, for example by fighting between males for a female, or by sneaking strategies to increase the number of matings [e.g. 4,5]. However, competition with other males can continue after mating, and can result in selection for increasing the number of male gametes transferred during mating, which increases the chance of fertilization [6–8]. Not only the *number* of gametes, but also their specific *characteristics* can increase the probability of achieving fertilizations. Sperm of animals often have adaptations that give them a competitive advantage in competition with other sperm [e.g. 9–11]. In plants, pollen are selected for increased growth speed of the pollen tubes to outcompete other pollen on the stigma [12,13].

Competition for mating occurs whenever there is a skew in the potential number of offspring produced by the two sexes. Generally, the sex producing the fewest but largest gametes, i.e. the females, is limiting for reproduction. This generally means that females can be choosy, while males are in competition for matings [14]. These two consequences of the difference between the sexes in investment in offspring can lead to a large variety of adaptations [2,14]. Only recently, it has been realised that these two aspects of sexual selection, viz. ‘male-male competition’ and ‘female choice’, also apply to fungi [15,16]. In many groups of fungi, different sex roles can be distinguished [17–19], which can result in different phenotypes [20], and there is a skew between the number of fertilizing individuals and individuals to be fertilized [21]. We have previously demonstrated that there is genetic variation in competitive ability and in choice between fungal individuals during mating [15]. So far, however, evidence for the occurrence of sexual selection in fungi is only circumstantial. In this paper, we directly test if sexual-selection theory, mostly developed for animals and plants, can also be applied to fungi, by performing an evolution experiment with the mushroom-forming basidiomycete *Schizophyllum commune*.

### *Schizophyllume commune* lifecycle

The general lifecycle of mushroom-forming basidiomycete fungi starts with the germination of a haploid meiotic spore, which grows vegetatively and forms a mycelium. This mycelium is called a monokaryon, as it contains haploid nuclei of only one type and in most species one per cell (see Fig 1). This mycelium can mate in a hermaphroditic fashion by hyphal fusion with a different mycelium. In its male role, the monokaryon fertilizes another mycelium by donating nuclei and in its female role, fertilizing nuclei are incorporated into the mycelium’s own cytoplasm [17]. Mating in basidiomycetes is only possible between monokaryons of different mating types. In most species, the mating type is defined by the alleles at the unlinked *A* and *B* loci, each composed of a cluster of tightly lined genes [22,23]. In *S. commune*, at each mating-type locus many different alleles exist [24] and as a result two monokaryons that meet at random have a very high chance to be compatible [25]. The fertilizing male nuclei are actively transported through the entire mycelium. Within the mycelium the migrating nuclei also divide and eventually the entire receiving mycelium becomes colonized by the fertilizing nuclei [26–28]. In the fertilized female mycelium, the male nuclei do not fuse with the resident female nuclei, but remain separate. This fertilized mycelium is thus entirely composed of cells with two genetically different haploid nuclei and is referred to as a dikaryon. After gamete fusion, nuclear fusion resulting in diploid cells is thus postponed and only occurs in highly specialized cells – the basidia–located in the sexual fruiting body – the mushroom, and is directly followed by meiosis. The meiotic spores formed on the basidia can disperse and germinate to start a new mycelium or can act as fertilizing propagules of an established monokaryon [29]. Because of the modular structure of the fungus, fruiting bodies can be formed anywhere on the mycelium. Fertilizing nuclei can therefore increase their fitness by occupying as much of the mycelium as fast as possible, before other nuclei colonize it [30,31]. In many basidiomycetes, nuclei migrate through the female mycelium at very high speeds, up to 90 times faster than mycelial growth rates [32,33].

**Figure 1.**
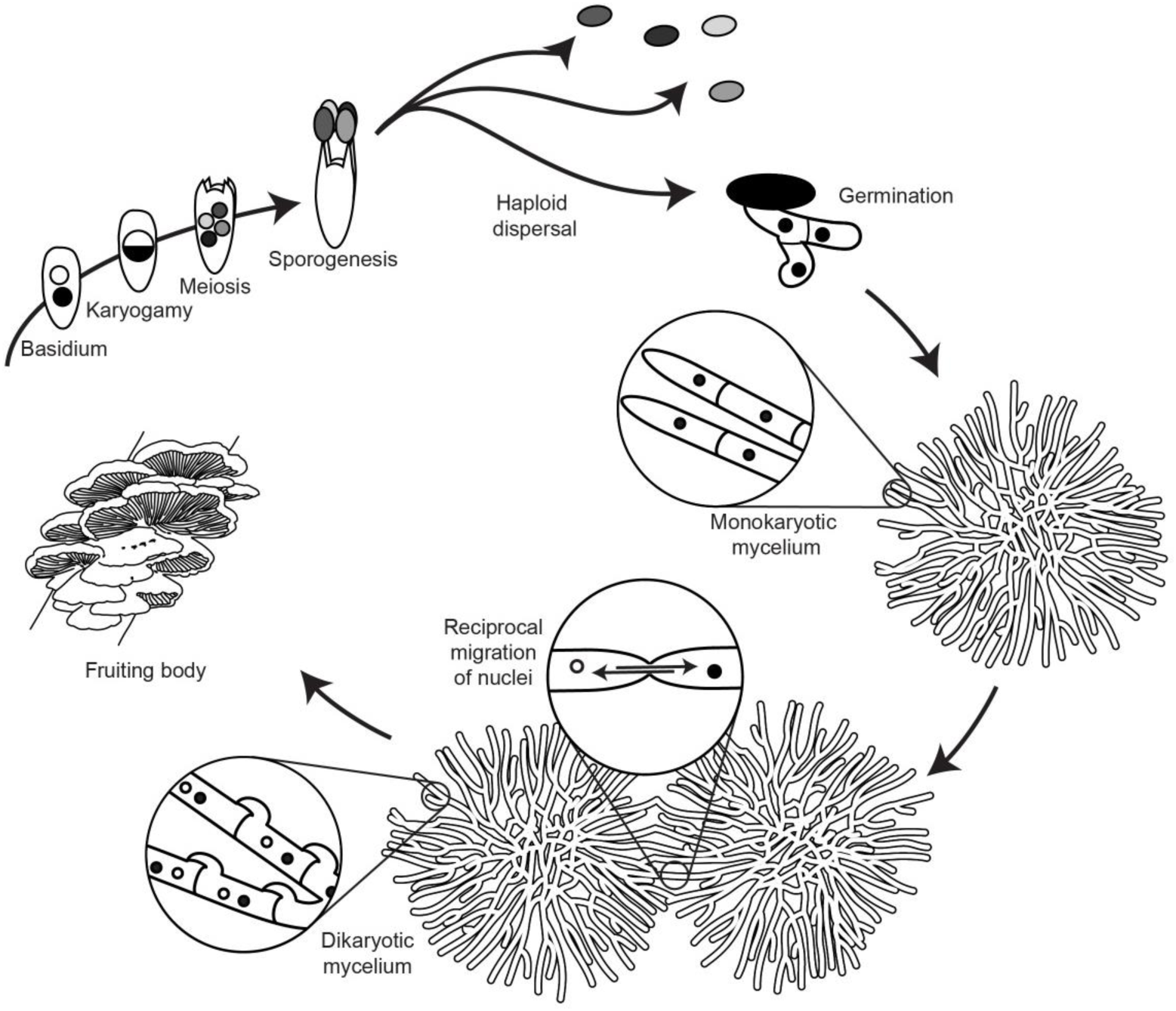
*Schizophyllum commune* lifecycle. A haploid spore germinates and grows by forming a mycelium - the monokaryon - in which a single type of haploid nuclei is present in all hyphae, and one per cell. Mating occurs by fusion of hyphae of different monokaryons, which exchange nuclei. The nuclei migrate from the point of plasmogamy and through the entire mycelium, thereby forming a long-lasting dikaryon, with two haploid nuclei per cell, one from each monokaryon. Fertilization can also occur unidirectionally by fusion with a spore or a dikaryon. When the conditions are suitable, the dikaryon forms a fruiting body, the mushroom, in which basidia are formed in which a short-lived diploid stage is immediately followed by meiosis and spore formation. The resulting haploid spores are wind-dispersed and can germinate to form a monokaryon.

We performed an evolution experiment with *S. commune*, in which an evolving population of nuclei was repeatedly allowed to mate in the male role with a non-evolving (‘naïve’) receiving female mycelium. We experimentally test the hypothesis that increased mating success in a mushroom-forming basidiomycete can be sexually selected, using an experimental evolution approach. We find that male competitive fitness after 20 sexual generations had significantly changed in two of twelve replicate lines, once increased, and once decreased. Competitive fitness was highly associated with colonization efficiency into the receiving mycelium, but not with nuclear migration rates. This suggests that a competitive advantage is obtained during the initial stages of entering the mycelium. Finally, we test the hypothesis that increased male fertility trades off with other fitness components, such as vegetative growth rate and fitness in the female role, and find that this generally is not the case.

## Material and Methods

### Outline of experimental setup

We performed an evolution experiment with the mushroom-forming basidiomycete *S. commune* in which an evolving population of nuclei was repeatedly mated in the male role with a non-evolving female monokaryotic mycelium (see Fig 2a). The nuclei migrated through the mycelium and fertilized it entirely. After the entire female mycelium was fertilized, the most distant part of the fertilized monokaryon – now a dikaryon – was transferred to a fresh plate and induced to form mushrooms and spores. All spores were then collected and used in a next cycle to fertilize a non-evolved unfertilized female mycelium. After 20 cycles of nuclear migration and sexual reproduction, the fitness of the evolved strains was measured, relative to the non-evolved parental strains. For the lines with a change in fitness, we also measured specific fitness components, which might have caused this change.

**Figure 2.**
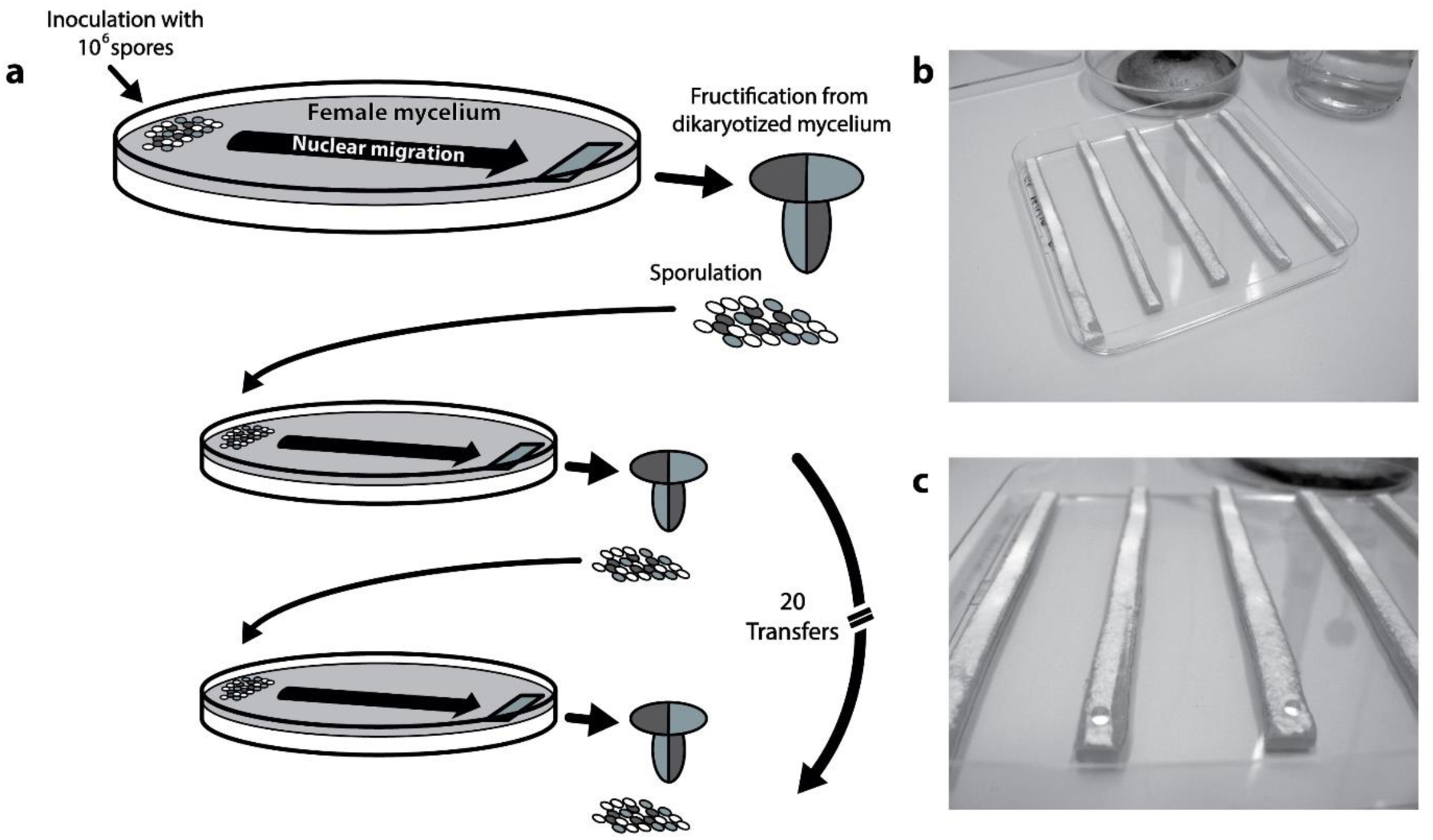
Setup evolution experiment and competition measurements. **a)** Basic setup of the evolution experiment. Spores (~10^6^) are inoculated onto a fresh receiving monokaryon. Spores can donate a nucleus to the mycelium, after which the nuclei can migrate through the mycelium for seven days. Next, a fragment of mycelium is removed from the far side of the plate and induced to produce mushrooms and sexual spores for 7 days. All spores produced by those mushrooms are harvested and used for a next round of selection. Each generation takes 14 days and we selected for 20 sexual generations. Dark grey symbolizes mating type A43B41 and light grey mating type A41B43. Spores are produced sexually and therefore half of the spores have parental mating types and half recombined mating types (white). Only 1/4 of the spores are compatible with the receiving monokaryon. All mycelia and spores contain the same resistance marker. **b-c)** Racetracks used for the competition experiments. On one side each 9cm long track was inoculated with a droplet containing a mixture of differently marked spores. A fragment of mycelium from the far end was taken after 7 days of migration and used to generate mushrooms of which the sexual spores were harvested and genotyped.

### Strains and culture conditions

*S. commune* strain H4-8 (matA43, matB41; FGSC no. 9210) [24] and H4-8b (matA41, matB43) [34] were used in this research (see Fig S1 for an overview of the strains and their names). A resistance marker against the antibiotic nourseothricin (N) or phleomycin (P) was introduced by transformation (plasmids and transformation protocol are described in [35]). The N marker was crossed into the compatible isogenic strain H4-8b. All strains used in this experiment were derived from a single cross between H4-8P and H4-8bN and are expected to be isogenic for all but mating type loci and resistance markers. Monokaryons are referred to as G1 or G2 for mating types A43B41 and A41B43, respectively and in addition, P or N to indicate resistance (e.g. G1P for a monokaryon with mating type A43B41 and phleomycin resistance marker). Dikaryons after 20 transfers are referred to by replicate line followed by marker type (for instance 3N for replicate line 3 with nourseothricin marker). Parental unevolved dikaryons are referred to as P0 (G1P x G2P) and N0 (G1N x G2N).

All culturing was performed on defined Minimal Medium (MM) [36] at 27°C in the dark. Mushrooms were formed in 12h dark 12h light cycles in vented 55mm plates, with Petri dishes placed upside-down, such that all spores produced would fall on the lid.

### Evolution protocol

For the evolution of increased fertilization capabilities, 20 replicate lines (ten of each resistance marker) were started, and were propagated using the following setup (see Fig 2). Mushrooms were produced from dikaryons P0 or N0. All spores were harvested in saline and a 40µl droplet containing ~10^6^ spores was inoculated on one side of a 9cm Petri dish covered with a female monokaryon. The female mycelium was grown for three days from macerated mycelium [15] from G1P or G1N for phleomycin or nourceothricin markers, respectively. The spores were allowed to fuse to the mycelium, after which their nuclei migrated through it and completely colonized it in 7 days. Even though two to three days is enough for a nucleus to reach the other side of the Petri dish, we waited longer to assure complete fertilization of the mycelium and increase the number of fertilizing nuclei. Seven days after inoculation with spores, a strip of agar (20×5 mm) with now dikaryotized mycelium was cut from the other side of the Petri dish, inoculated on a fresh plate and grown for seven days in dark-light (12h-12h). During this time the dikaryotic mycelium grew vegetatively and produced dozens of tiny mushrooms, which produced spores. All spores were harvested in 1ml saline. This suspension was centrifuged for 5 min at 3800 *g* after which the bottom most 40 µl again was used to inoculate a fresh unevolved monokaryon with the corresponding resistance marker. Even though during evolution one of the lines (6N) started to produce an increased number of spores, more than fit in 40 µl, always only spores from the bottom 40 µl were used. This procedure was repeated for 20 sexual generations.

For future reference, 2×2mm blocks of agar with mycelium were cut out of the dikaryon that grows next to the mushrooms and immediately frozen at −80°C. This is a convenient way of storing the mycelium, because a single small block from −80°C can be placed immediately on a fresh MM plate and will continue to grow. Additionally, after every fifth transfer dikaryotic mycelium was grown on cellophane and stored in 15% glycerol at −80°C.

### Competitive fitness measurements

Competitive fitness assays were performed in a setup similar to the selection regime. The relative fitness of evolved lines at transfer 20, was measured in direct competition with the parental strains. Equal numbers of spores from parental and evolved strains with different markers were inoculated onto an unevolved monokaryon, one round of selection was performed, and the relative contribution to the next generation was measured. To obtain spores, mushrooms were grown from mycelium revived from the −80°C. An agar block of to-be-revived dikaryons stored at −80°C was placed on fresh MM and grown for 1 week. From this plate a 5×20 mm piece of agar was taken from which mushrooms were grown.

Fitness could not be measured for eight of the 20 selection lines. Lines 5P and 10N stopped producing mushrooms during evolution (after 15 and 17 transfers respectively), even though dikaryons with clamp connections had been established. Additionally, six lines (5N, 1P, 4P, 6P, 7P, 8P) were incapable of producing mushrooms after reviving the evolved strain from the −80°C, even though after the last transfer mushrooms from the same mycelium were present. Even though more P than N strains did not fructify (6 and 2 respectively), this does not deviate from expected effect (Fisher exact test, *p* = 0.170). We were not able to induce fructification in these lines using a variety of standard manipulations [37], therefore neither fitness assays nor follow-up investigations could be conducted on them. For the remaining 12 evolved lines fitness was measured.

To obtain enough spores for all competitions, 20 plates were inoculated for parental lines, ten of each marker, from which all spores were collected and pooled. All spores were harvested and suspended in 1 ml saline. This suspension was centrifuged (5 min at 3800 *g*) after which 900µl supernatant was removed to concentrate the spores. Next, the spore density of this suspension was determined by counting with a hemocytometer, and the suspension was diluted to 10^7^ spores/ml. For each evolved line, 100µl of this suspension was mixed with an equal volume with the same concentration of spore suspension of the parental strain that carried the alternative resistance marker and vortexed twice, to obtain a homogeneous dilution of 10^7^ spores/ml with equal numbers of parental and evolved spores. Of this mixture, 10µl was pipetted on one side of a ‘competition track’ of unmated mycelium without a resistance marker. A ‘competition track’ was created by cutting strips of agar (90×8mm) covered with mycelium of monokaryon G1 (i.e. without resistance marker) from a square 12cm Petri dish with 45ml MM. Five tracks were lined next to each other in an empty 12cm square Petri dish (see Fig 2b). In every second plate one track was not inoculated to check for cross fertilizations, which were never observed. To assess the exact frequency of evolved and parental strain in the inoculum (quantitative PCR – see below – was not possible due to low concentrations), three dilutions of the mixture were plated on MM+0.5ml 5% Triton-80 to form colonies in six replicates. After three days the colonies of those dilutions that yielded between 50 and 500 colonies were counted, after which each plate was covered with 500µl antibiotic of either nourseothricin (0.4 mg/ml) or phleomycin (1.25 mg/ml). After two days the colonies that continued to grow were counted. Per line nine replicate measurements were performed. Additionally, the parental strains were competed against each other for control.

After 7 days, the last 5mm of the track was placed on a fresh MM plate to form mushrooms and spores. To determine the frequency of the evolved vs. parental types we performed quantitative PCR using the resistance genes as targets (NFw: 5’-CACTCTTGACGACACGGCTTAC, NRev: 5-AAGGACCCATCCAGTGCCTC, PFw: 5’-AAGTTGACCAGTGCCGTTCC and PRev: 5’-AAGTCGTCCTCCACGAAGT). The spores produced are the result of a meiosis from a fusion of one nucleus from the unmarked receiving mycelium, and one from either of the two marked fertilizing spores. Therefore, half of the spores will not carry any marker and the other half can be of either marker, following Mendelian segregation. If both nuclei perform equally well during fertilization, the ratio of the markers is 1:1 and if one performs better, a deviation from 1:1 will be observed. Using colonies from after selection would be too cumbersome due to the large number of replicates per strain. Controls using both colonies and qPCR showed equal ratios for both methods (see Fig S2). The qPCR method was tested and optimized using a monokaryotic strain carrying both markers as single copy inserts, which made is possible to measure the efficiency of the qPCR reaction for each marker at exactly equal frequencies. These efficiencies were used to calculate the frequencies of each marker within one sample [38].

Spores were collected for DNA isolation to be used in qPCR. All spores were harvested in 700µl LETS buffer and spore walls were destroyed by freezing the dilution at −20°C followed by incubation with 20µl Proteinase K (20 mg/ml) for 4h at 56°C, after which standard phenol/chloroform extraction was performed [39]. Each reaction consisted of 5µl of undiluted DNA and 5 µl SYBR Green 2X (Biorad) with 200 nM final concentration of each primer and was performed on a Biorad CFX96 machine. For each sample, all reactions for both loci were performed in the same run with one technical replicate per locus. The cycling conditions were 1 cycle at 95°C/10 min, followed by 40 cycles of amplification (95°C/10 sec, 62.5°C/30 sec, followed by a fluorescence read). For analysis we used the software package CFX Manager (Biorad) with standard settings for baseline and thresholds. The ratios of marked strains (R = Evolved/Parental) before and after competition were used to estimate the relative competitive fitness (fitness = Log_10_[R_before_/R_after_]) [40].

### Fitness components

For five evolved lines that showed changed competitive fitness before correction for multiple measurements (6N, 7N, 2P, 9P and 10P; see ‘Results’) and the parental strains (P0 and N0), specific components of fitness were measured which we inferred played a role during selection: 1) spore production and size, 2) spore germination, 3) mating type of the spores, 4) nuclear migration speed, 5) establishment potential in the female mycelium (which we termed ‘colonization efficiency’), 6) growth as a dikaryon, and 7) mushroom formation.

#### Spores: yield, size, and germination rate

The total number of spores produced seven days after inoculation of the dikaryon, were measured on a Z2 Coulter counter (BeckmanCoulter), for three biological replicates each. Spore size was measured by taking photographs at 200X magnification with a phase contrast microscope and measuring the length of at least 40 spores for three replicates each using ImageJ V1.44n [41]. To assure the spores had not started germinating, spores that were produced within 2 hours before measurement were harvested in saline and immediately stored on ice.

Germination was measured after spores were concentrated in saline solution, mimicking circumstances in the evolution experiment. The spores were harvested in saline, spun down, and the bottom 40 µl solution was transferred to fresh 1.5 ml reaction tubes with small holes for air exchange and allowed for germination. After 24 h, 48 h and 72 h the number of germinated spores of a total of 100 spores was counted at 200X magnification in a blind setup. Per line three biological replicates were performed.

#### Mating-type ratio

Because only spores of mating type A41B43 are compatible with monokaryon G1 (A43B41), selection for increased proportion of A41 or B43 alleles might have occurred during evolution. Alternatively, linkage between the two mating-type loci can increase compatibility potential two-fold [25,42,43]. For each line at least 48 spores were isolated, and their mating type was determined by performing crosses following Papazian [44].

#### Nuclear migration speed

Per line, 20 ‘competition tracks’ (see above) of 12 cm length were inoculated with ~10^6^ spores on one side and incubated. Each 24h, five random racetracks per line were sacrificed and cut into 1 cm pieces. Each piece was placed in a well of a 24 wells plate containing 0.5ml MM and incubated for two days. After two days outgrowth, each piece was checked for clamp connections to determine if the piece of mycelium had become dikaryotized, which indicates that at least one fertilizing nucleus was present in that piece of mycelium at the time of isolation [26].

#### Colonization efficiency

The competitive ability for colonization of the female mycelium after 7 days was measured by measuring the abundance of fertilizing nuclei relative to the parental strain. For the five lines with fitness change and the parental strains against each other, we repeated the measurements as described in ‘Competitive fitness measurements’, and additionally, we sampled the mycelium in the ‘competition track’ at two points, 10 and 50 mm after inoculation. At each position 10mm of the track was cut out from which DNA was isolated using QIAGEN DNEasy Plant mini kit. qPCR was performed on undiluted DNA derived from spores and 10 times diluted DNA from the mycelium. Relative competitive colonization efficiency was calculated as described above. This experiment was performed with two different female mycelia: nine replicates with monokaryon G1 as receiver, and nine replicates with monokaryon G2 as receiver.

#### Mycelium growth rate of monokaryon and dikaryon

Finally, from each dikaryon, 24 single spore colonies were isolated. For each monokaryotic isolate a small mycelium plug was inoculated on a 9cm Petri dish and radial growth was measured after 3 days of incubation. The same was done in threefold for each dikaryon.

### Statistical analyses

For the relative fitness measures and colonisation efficiency comparisons, the means of the evolved lines relative to means of the parental strain were compared using a two-tailed Dunnett’s test. The same test was used for spore size and spore yield. For germination speed, nuclear migration and mycelial growth, Kruskal-Wallis tests were performed with pairwise comparison Wilcoxon test with Bonferroni correction as post-hoc test. Mendelian segregation of mating types for all strains was tested using Pearson’s χ^2^ tests. All statistical analyses were performed with R 3.0.1 [45].

## Results

Experimentally evolved strains adapted to fertilize a non-evolving female mycelium were analysed for competitive fitness and components of fitness. First, the results for fitness are given, then the components in the order as occurring during fertilization.

### Competitive fitness

Fitness was measured relative to the parental strain. From the twelve evolved lines for which we could measure fitness (see ‘Material and Methods’ for explanation on the other 8 lines), five had significantly changed fitness (see Fig 3, Student’s T-test, p < 0.05). Four lines (6N, 2P, 9P and 10P) showed an increase in fitness relative to the parental strain and one (7N) a decrease. The two parental strains showed no marker effect (fitness_N0 relative to P0_ = 0.005, S.E. = 0.206, p>0.1). A Dunnett’s test showed that only 6N and 7N are significantly different from the parental when correcting for multiple comparisons. Further investigations of the five strains with (marginally) changed fitness showed that the found differences were robust, except for strain 2P.

**Figure 3.**
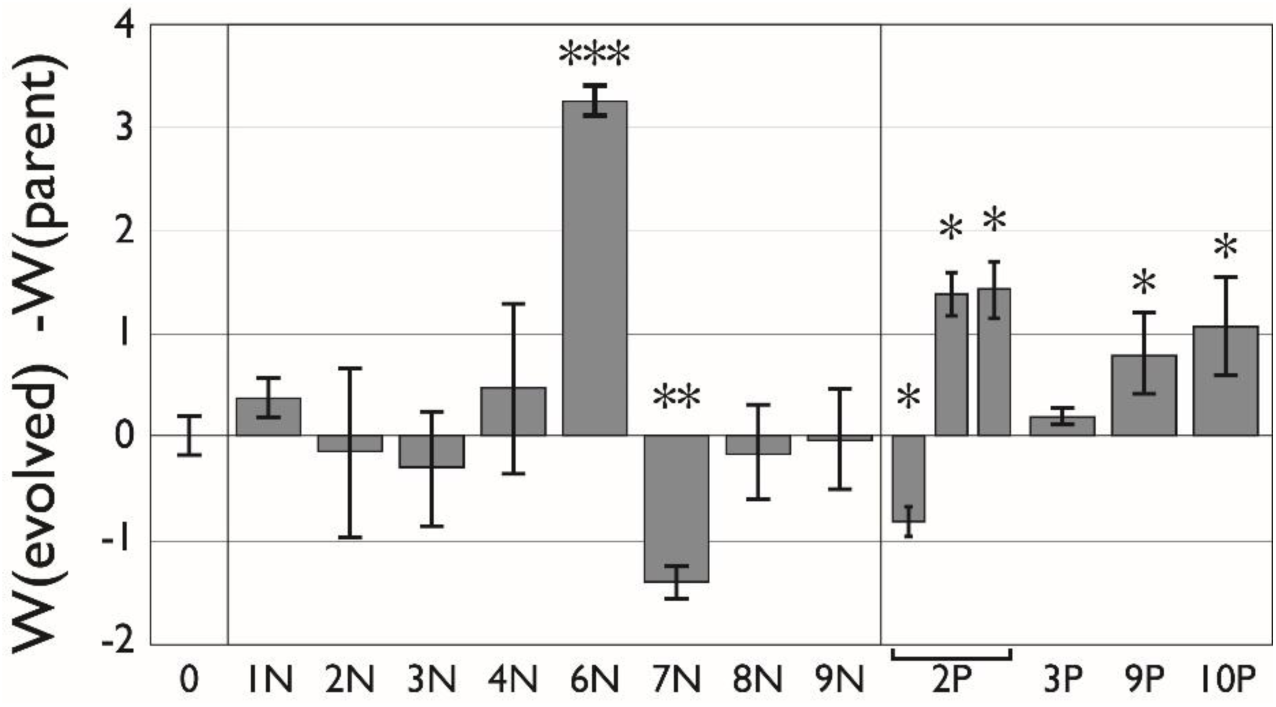
Relative fitness of the 12 tested strains. 0 indicates competition between the parental strains P0 & N0. The other values are competitions of lines from transfer 20, relative to the parental strain of the opposite resistance marker. Positive values indicate increased fitness and negative reduced fitness. Measurements of 9 replicates. Error bars indicate standard error. (*: *p* < 0.05, **: *p* < 0.01, ***: *p* < 0.001).

Competition with line 2P in the initial fitness measurements showed a reduction in fitness (W_2P_ = –0.732, S.E. = 0.324). When testing competitive fitness with the ‘colonization efficiency’ however we found an *increase* in fitness both for G1 and G2 (W_2P_ = 1.404, S.E. = 0.291 and W_2P_ = 1.900, S.E. = 0.363 respectively). All the other lines showed similar results between both replicates. We therefore performed a third assay in which 2P again was tested in competition with N0 which showed results similar to the ‘colonization efficiency’ measures (W_2P_ = 1.391, S.E. = 0.227). It is unclear why the results in the first assay differed so much from the others. In Fig 3, data from all three assays are presented.

In competitions with line 6N the parental strain was below detection levels (i.e. 40 PCR cycles) and to be able to calculate a fitness value the parental strain was set to 40 cycles. The fitness value of line 6N should thus be considered as a minimal fitness measure. Tests between strain 6N and 9P – the fittest evolved phleomycin strain – yielded similar results (data not shown).

### Fitness components

Six different fitness components were measured for the parental strain and the five lines that showed (marginal) fitness change. Characteristics that are expected to be of importance during mating, as well as vegetative characteristics, which may have changed as pleiotropic consequences of increased competitive fitness, were measured.

#### Spore characteristics

##### Spore size

Spores collected from mushrooms produced by evolved lines showed no difference in spore size or shape relative to unevolved parental strains (data not shown).

##### Germination speed

Germination was measured 48 and 72 hours after incubation at 27°C (Fig 4a). Line 2P had increased spore germination at 72h. Line 7N had strongly reduced spore germination after 72h compared to the parental strain (8.7% and 37.3% for 7N and parent respectively; Bonferroni corrected two-sample t-tests, p < 0.0083). Measuring germination at later time points became impossible, due to mycelial growth of previously germinated spores.

**Figure 4.**
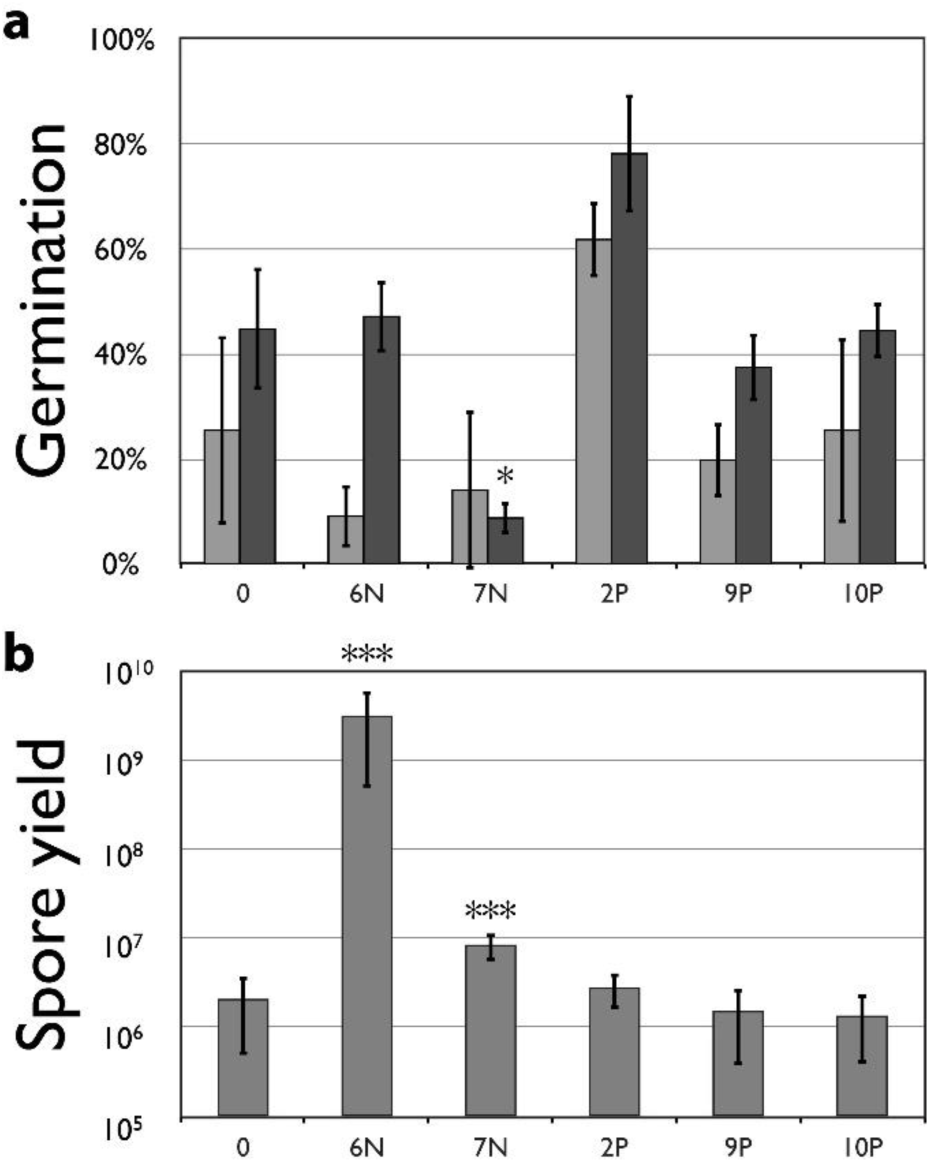
**a)** Spore germination percentage after 48 hours (light grey) and 72 hours (dark grey). **b)** Average spore yield. Total number of spores produced after 7 days for parental and evolved strains at transfer 20. Y-axis is log scaled. (Error bars indicate standard error; Significance values are relative to values from the parental strain. *: *p* < 0.05, ***: *p* < 0.001).

##### Spore yield

For two lines – line 6N and 7N – spore yield had increased relative to the parental strain (Kruskal-Wallis, *p* < 0.0001, Fig 4b). Line 7N, which had reduced competitive fitness relative to the parental strain (see Fig 3), showed a more than 4-fold increase in spore production. Nevertheless, this line had a decrease in competitive fitness. Line 6N had a stunning 1500-fold increase in spore production relative to the parental strain. The increase in spore number in line 6N started already after four transfers. Associated with this increased spore production, 6N showed a changed morphology of the fruiting body. At the fourth transfer during the evolution phase, part of the mycelium did not produce many small mushrooms as seen in the parental strain (Fig 5a), but produced larger mushrooms directly on the mycelium (Fig 5b-f) and from transfer 9, the line appeared to produce one single large mushroom. This mushroom phenotype produced many more spores and is likely a reversion to the level of spore production seen in natural isolates. Mushrooms produced by dikaryons consisting of a nucleus derived from natural isolates and one either from the parental monokaryons, or from monokaryotic offspring from 6N, all produced spores in quantities comparable to 6N. Therefore, low spore production apparently is a recessive lab strain specific characteristic. Analysis of monokaryotic offspring from line 6N showed that the mutation causing increased spore production segregates with the A mating type locus, showing that there is genetic linkage (5.5 cM) with the A locus (Fig 5g).

**Figure 5.**
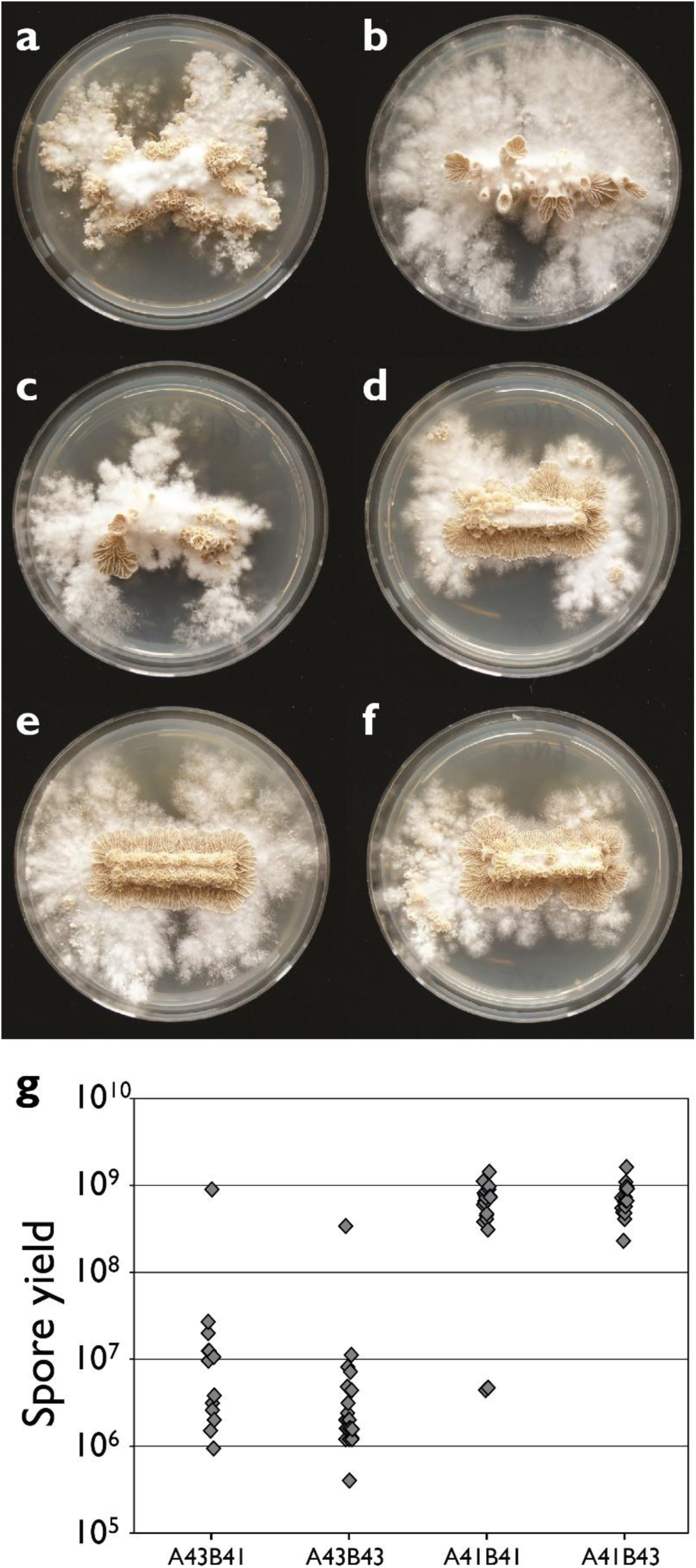
Mushroom morphology and spore production. Change in mushroom morphology of line 6N. After 0, 4, 8, 10, 15 and 20 (**a** to **f** respectively) sexual generations of experimental evolution. Pictures show mushrooms grown on 5cm plates. To obtain mushrooms, dikaryotic mycelium was revived from the −80°C, grown for 5 days to completely cover a 55mm plate. Then a piece of approximately 20 × 5 mm was transferred to a fresh plate which was incubated for 7 days with the plates placed upside-down. **g**) A qtl for spore production is found linked to the A mating-type locus.

#### Nuclear migration speed

All evolved strains were able to fertilize the mycelium and no differences were observed between the parental strains and any of the evolved lines (Kruskal-Wallis, p > 0.05; Fig S3). After 24 hours, migration had occurred in most replicates and average migration was 16.8 mm which increased to 75.2 mm after 48h and 114.4 mm after 72h. On day 4, the final centimetre of the tracks was reached in all replicates for all lines.

#### Colonization efficiency

We measured the ratios of fertilizing nuclei after seven days in the female monokaryons G1 and G2 (Figs 6a and 6b respectively), 10mm and 50mm from the point of inoculation, as well as the overall fitness (*i.e.* the representation of the evolved and parental genomes in the spores formed, as previously described). Measurements of nuclear ratios in the mycelium showed that already after 10mm the ratio between the two nuclei was highly skewed, and that these values could almost completely explain the final fitness increase measured after spore formation. Line 7N shows a negative value, indicating that the loss in fitness might be caused by a change in the same mechanism that gives the other lines an advantage. These findings suggest that the competitive advantage of the evolved lines was obtained prior to or within the first 10mm of migration. This concurs with the findings by Ellingboe [46] on nuclear migration in dikaryon-monokaryon matings, in which a nucleus successful in establishing in the mycelium migrates with the same speed, but is ahead in the advancing front. Except for 10P, no significant further increase in nucleus ratio was observed between the measurement at 10mm and later.

**Figure 6.**
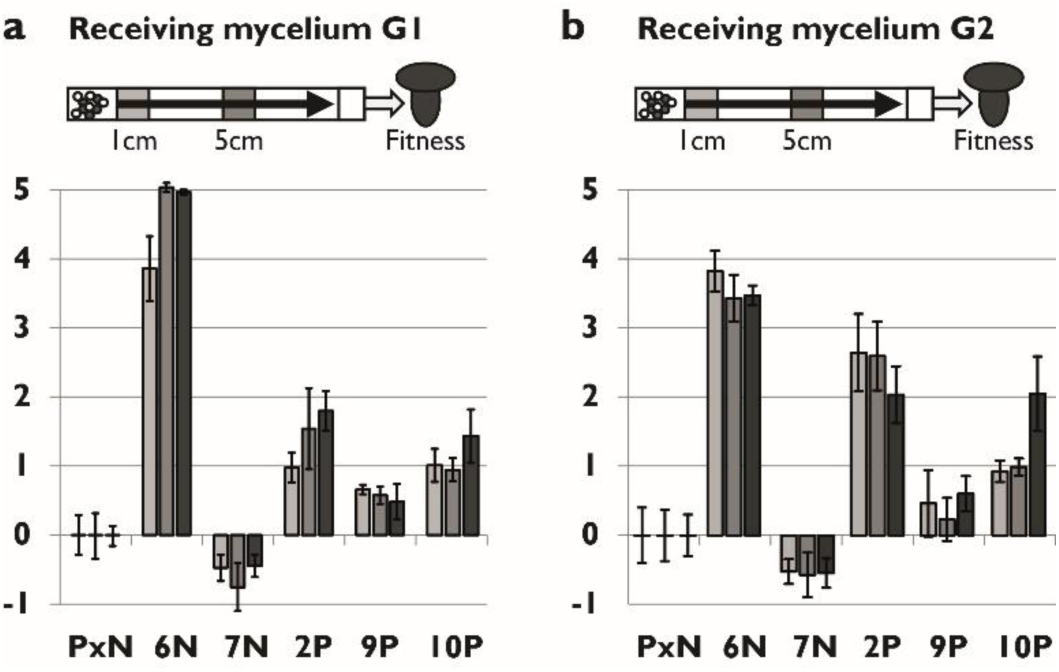
Colonization efficiency and relative competitive fitness of the parental line and the five evolved lines with changed fitness. **a)** Bars indicate relative competitive ability in colonization of mycelium G1 (mycelium used as female during evolution) after 1 and 5 cm migration, as indicated in the figure above each graph (first two bars) and relative competitive fitness (light bar). **b)** as a) but with receiving mycelium G2 used. Measurements of 9 replicates. Error bars are standard error.

Strikingly, all lines except for 9P, had similar fitness when mated to the G1 or the G2 strain. The first is the female mycelium to which adaptation could have occurred during evolution, while the latter is an isogamic strain of the opposite mating type.

#### Mating-type ratios

During meiosis, Mendelian segregation of the mating-type alleles will yield spores of four different genotypes (A43B41, A43B43, A41B41 and A41B43). Only spores of type A41B43 are compatible with the receiving mycelium G1 (A43B41). Theoretically, there are several possibilities for sexual selection to increase the fraction of compatible spores. First, meiotic drive for either the A41 or B43 allele will produce more compatible spores. Second, linkage (or a modifier for repressed recombination) between mating-type alleles A41 and B43 would increase the fraction of compatible spores from 25% to 50% [47]. To explore these possibilities, single spore cultures were isolated from the evolved lines, and their mating types were determined (Table 2). No deviation from Mendelian expectations was detected (χ^2^-test, α = 0.05, df = 3).

**Table 1.** Overview of fitness and migration ratio for evolved.

| Strain | Relative fitness<sup>a</sup> |  | Mating to G1 |  |  | Mating to G2 |  |  |
| --- | --- | --- | --- | --- | --- | --- | --- | --- |
|  |  |  | 1 cm | 5 cm | Spores | 1 cm | 5 cm | Spores |
| **0** | 0.00±0.08 | - | 0.0±0.30 | 0.0±0.33 | 0.0±0.15 | 0.0±0.40 | 0.0±0.37 | 0.0±0.30 |
| **1N** | 0.35±0.16 | 0.0666 | – | – | – | – | – | – |
| **2N** | -0.2±0.37 | 0.5981 | – | – | – | – | – | – |
| **3N** | -0.08±0.50 | 0.8807 | – | – | – | – | – | – |
| **4N** | 0.33±0.35 | 0.3708 | – | – | – | – | – | – |
| **6N** | 3.23±0.15 | <0.001 | 3.86±0.47 | 5.04±0.07 | 4.97±0.04 | 3.81±0.30 | 3.42±0.33 | 3.46±0.14 |
| **7N** | -1.31±0.16 | <0.001 | -0.47±0.19 | -0.75±0.34 | -0.45±0.16 | -0.52±0.18 | -0.57±0.32 | 0.01±0.21 |
| **8N** | -0.06±0.45 | 0.9004 | – | – | – | – | – | – |
| **9N** | 0.07±0.48 | 0.8879 | – | – | – | – | – | – |
| **2P** | -0.71±0.27 | 0.0425 | 0.98±0.22 | 1.54±0.59 | 1.8±0.29 | 2.63±0.55 | 2.6±0.50 | 2.03±0.41 |
| **3P** | 0.09±0.17 | 0.6527 | – | – | – | – | – | – |
| **9P** | 1.20±0.30 | 0.0064 | 0.66±0.07 | 0.58±0.12 | 0.48±0.26 | 0.46±0.48 | 0.22±0.31 | 0.40±0.25 |
| **10P** | 1.33±0.26 | 0.0011 | 1.01±0.24 | 0.95±0.17 | 1.43±0.39 | 0.92±0.15 | 0.99±0.12 | 2.04±0.54 |

<sup>a</sup> Mean relative fitness ± standard error and *p* value for two paired t-test with parental strain.

**Table 2.** Mating type percentage of *n* tested single spore isolates derived from parental and evolved lines. A43B41 was used as the female mycelium (monokaryon G1) during evolution, A41B43 the compatible male mating type.

| Strain | A43B41<br>♀ | A43B43 | A41B41 | A41B43<br>♂ | $n$ | $\chi^2$ |
| --- | --- | --- | --- | --- | --- | --- |
| 0 | 25.0% | 20.8% | 33.3% | 20.8% | 48 | 2.00 |
| 6N | 18.8% | 29.0% | 29.0% | 23.2% | 69 | 2.01 |
| 7N | 21.6% | 21.6% | 28.4% | 28.4% | 116 | 2.21 |
| 2P | 27.1% | 22.9% | 29.2% | 20.8% | 48 | 2.17 |
| 9P | 29.2% | 20.8% | 18.8% | 31.3% | 48 | 0.83 |
| 10P | 22.9% | 27.1% | 20.0% | 30.0% | 70 | 0.17 |

#### Mycelium growth rate

##### Dikaryon growth rate

Growth rate of the evolved lines was measured over three days of growth, for which we used the fertilized mycelium from generation 20, a dikaryon composed of the resident female nucleus and probably a population of fertilizing male nuclei. Only the dikaryon 10P had a changed growth rate which was strongly reduced relative to the parental strain (p < 0.001, 3.46 mm/day, S.E. = 0.054). No difference was seen for the other dikaryons (p > 0.05, mean growth = 8.96 mm/day, S.E. = 0.273).

##### Monokaryon growth rate

For the monokaryons derived from each evolved lines, growth rate was increased for 2P and decreased for 9P (see Fig 7a, post-hoc Bonferroni corrected Wilcoxon rank sum test, p = 0.005 and p = 0.003 respectively). 2P shows an increased growth rate on average, but with a similar distribution as seen in the parental strain (Fig 7b-c). 9P shows a bimodal distribution in which 10 strains have reduced growth rate (Wilcox Rank Sum test, p < 0.001) and 14 a growth rate similar to that of the parent (Fig 7d). Growth rate is not associated with either mating type (Wilcox Rank Sum test, p > 0.5).

**Figure 7.**
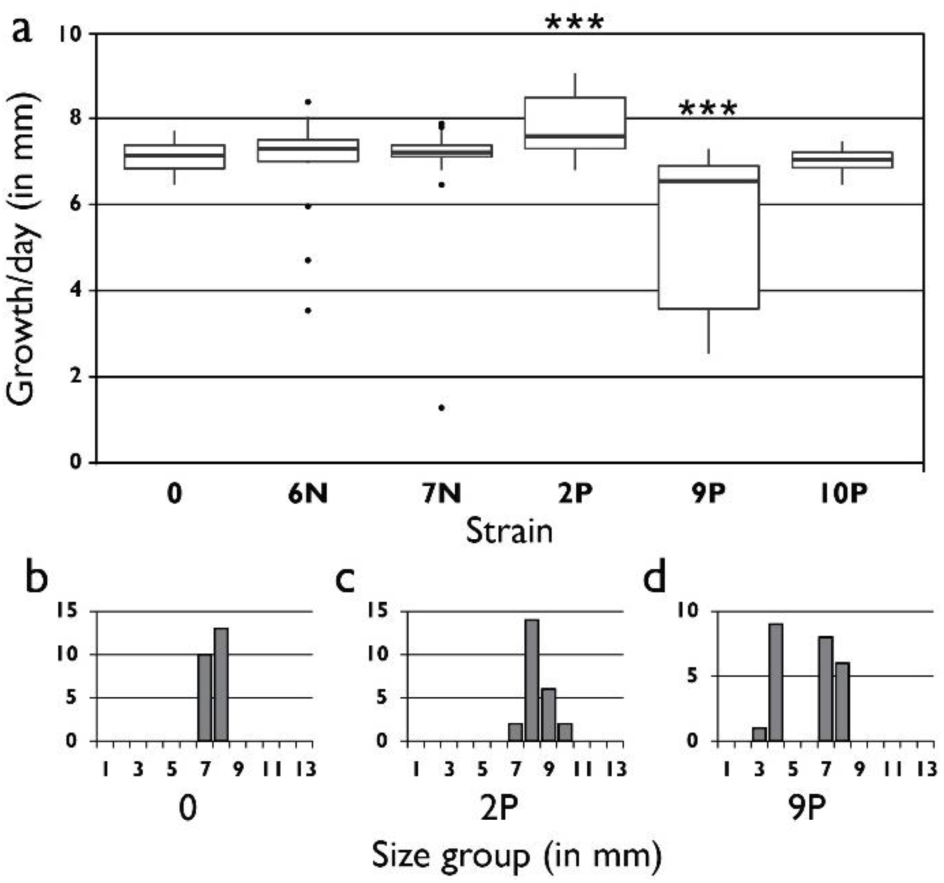
Monokaryon mycelium growth rate. **a)** Box plot of growth rate of 24 monokaryons derived from parental and evolved strains at transfer 20 (**: p < 0.01). **b-d)** Histogram of data per strain for parental strain and the two evolved strains with changed monokaryon growth rate. Strain 9P shows a clear bimodal distribution with a slow and a fast growing group. The mycelia from the fast growing group grew with the same rate as the parental strain.

## Discussion

We have experimentally shown that there is potential for the evolution of increased mating success in a mushroom-forming basidiomycete. By repeatedly mating a monokaryon with a non-evolving receiving ‘female’ monokaryon, two out of twelve lines showed a significant change in reproductive success: one with increased, and one with decreased relative fitness. A further two lines showed a change that after correction for multiple measures was not significant at the 0.05 level, but in subsequent measures showed reproducible results, providing circumstantial evidence for fitness change in those lines as well. In a fifth line inconsistent fitness measures were found. For these five lines, we measured a variety of mating components to study which traits are responsible for changes in competitive fitness, and found a variety of traits that changed relative to the unevolved controls. We found that the main determinant of success in a mating is the nuclear ratio in the mycelium, which already after 10mm of nuclear migration is indicative for the final fitness. This ratio is most likely caused by the speed of basidiospore germination. An additional adaptation is the total number of spores produced, which likely directly increases fitness. We did not observe direct trade-offs between fertilization success as males and female or vegetative traits. Our findings give insights into those components that influence the functionality of basidiospores as fertilizing units [18,48] and the male mating success of a fertilizing nucleus [16,23].

One unexpected finding was a consistent *decrease* in fitness in replicate line 7N relative to the parental strain after 20 transfers. Experimental evolution in large populations is expected to result in increased fitness, not in a decrease [49]. One possible explanation for the observed reduction in fitness is that a deleterious mutation became fixed due to genetic drift caused by a small effective population size of selected nuclei. Indeed, on some occasions some strains produced only few basidiospores at the moment of transfer. This is further supported by the loss of sexual reproduction in two of the strains during our experiment. Nevertheless, deleterious mutations are not expected to be maintained. Each round, the genome of the migrating nucleus recombines with the parental strain, during which the new deleterious mutation can be purged. Only when a mutation is located in or closely linked to one of the two selected mating type alleles – the only regions that cannot be ‘refreshed’ from the female mycelium – will a deleterious mutation be maintained too. The reduced fitness observed in line 7N might thus have been caused by fixation of a dominant deleterious mutation in or tightly linked to either the matA41 or matB43 allele. Most likely, fitness is decreased due to reduced germination rate of the spores (see Fig 4a). That the mutation influences fitness in the dikaryotic or diploid stage and is dominant can be deduced from the observation that fitness is reduced both in matings with G1 and G2 (see Fig 6). Even though the line did show increased spore production (Fig 4b), this did not compensate for the overall fitness reduction.

### Selection for mating success

#### The number of gametes

An important component in obtaining matings and fertilizations is the sheer number of – generally male – gametes that are interacting in the competition for obtaining gametes of the other sex – generally female. Bateman [50] suggested that when a female mates with multiple males, a male with more sperm will increase its success in mating. This was later formalized by Parker and others [6,51] and much empirical evidence has been generated that supports these findings [52,53]. Also in plants increased pollen production leads to higher mating success [54,55].

Two of the lines in our selection experiment with changed competitive fitness (6N and 7N) had increased spore production. Line 6N had the highest fitness increase and also had a more than 1000-fold increase in spore yield, but line 7N, which produced four times more spores than the unevolved lines, had decreased fitness. If competition is a fair raffle in which each gamete has the same chance of fertilizing, fitness would be proportional to gamete frequency. The results from these two lines suggest that the sheer number of spores produced is not the only determinant of mating success. First, even though line 7N had increased spore production, it had a reduced fitness, and second, line 6N had increased fertilization successes in the female mycelium, even when the inoculum for the competitions was controlled to a 1:1 ratio between evolved and parental spores. Even though higher spore production is likely beneficial during mating – because we transferred all spores produced, this increases fertilization success in a fair raffle – (see below), gamete quality is probably more important. The high number of spores produced additionally could have aided adaptation in these lines by increasing the effective population size and hence the mutation supply rate.

#### Characteristics of gametes

The importance of gamete quality in animals (e.g. sperm velocity [53,56,57]) and plants (e.g. pollen size[58]) on fertilization success has been well documented. Quality of the gametes seems to be an important trait in our experimentally evolved lines too. The ratio of parental and evolved fertilizing nuclei shortly after initial contact with the female mycelium, was skewed in the same direction as fitness and remained stable during migration (see Fig 6 and Table 1). Fast access to the mycelium might help establish this dominance, similar to sperm precedence. A better analogy might be with pollination in plants, where pollen that germinate faster [59], can access the stigma faster (sometimes by manipulation of the female stigma [60]). Also, pollen with faster tube growth [13] have increased fertilization success.

#### Spore germination

The first step to increase fertilization success is by increased access to the female mycelium by faster germination or increased germination success. Indeed, line 2P showed increased germination rate and germination success (see Fig 4a), which might give this line an advantage in fertilizing the monokaryon. Line 7N showed reduced germination success relative to the parental strain, which might explain the observed reduction in fitness for this strain. No difference in germination was observed for the other lines with increased fitness.

#### Fusion to the female mycelium

The second step to increase fertilization success is by improving fusion to the female mycelium, either directly between spores and mycelium [61] or directly after germination between the two monokaryons. If a basidiospore of *S. commune* is close enough (<~15 µm) to the female mycelium it can attract female hyphae (similar to the trichogyne in ascomycetes), which grow towards it and fuse, probably by producing a chemical used for chemotaxis by the hypha [61] [62]. If spores produce more of this chemical, they might be more successful in attracting hyphae and increase their head start in migration. Obvious candidates for such a compound are the pheromones encoded on the *B* mating type which are used for extracellular communication [63,64]. If extracellular communication changed attractiveness, it is unlikely that the pheromones themselves changed, because fitness was similarly affected in matings with either G1 or G2. An alternative possibility is that the amount of pheromones secreted is increased post-transcriptionally or that there is a change in secretion machinery. Increased pheromone production has been shown to increase mating success in *Saccharomyces cerevisiae* [65,66].

#### Migration rate

The third step to increase fertilization success is by faster nuclear migration. The migration rate of nuclei in basidiomycete fungi has an analogy with sperm velocity in animals [67] and pollen-tube growth in plants [13,68]. Increased migration rate would be advantageous for male function, because it increases colonization speed, which could give a competitive benefit if the colonized parts of the mycelium are closed to later arriving nuclei. However, as observed by Ellingboe [46] in dikaryon-monokaryon matings, genetically different nuclei that migrate into a mycelium to fertilize it, travel with approximately the same speed. Similar to Ellingboe, we did not measure significant differences in migration speed between evolved and ancestral strains. Furthermore, similar to our observations, Ellingboe observed that the nucleus eventually winning the fertilization in these experiments was the one that established in the mycelium more efficiently, thereby gaining a head start.

It is well possible that nuclear migration speed is determined and limited by the receiving mycelium, which reduces the potential for adaptation in this trait [28] and would explain that in our selection experiment, migration speed did not change. Furthermore, contrary to plants and animals, high migration rates are probably beneficial for female function too. High migration rates increase the chance that a monokaryon completely becomes fertilized quickly after mating, thereby avoiding additional fertilizations which would result in the split of the female mycelium into two or more different dikaryons. For many plant and animal species, multiple matings can increase female fitness if it increases male-male competition, which has led to selection for increased obstacles for male gametes [69–71]. In basidiomycetes, however, balkanization of the mycelium is likely to reduce total fitness due to negative interaction between the different dikaryons [72,73]. A single mating would maximize fitness for the female mycelium and thus select for mycelia that increase fertilization speed.

### Male and female roles during mating

Theory predicts that male and female reproductive success in hermaphrodites trade off in some form [74,75]. Even though male and female sex roles in a mushroom fungus cannot unambiguously be assigned to all aspects of the reproductive cycle – for example the dikaryon phase is not associated with either sex – in general, male traits are associated with fertilization, and female traits with becoming fertilized [21]. Exclusively female traits are acceptance of male nuclei into the own mycelium and monokaryotic growth, although the latter might also benefit male function because it can increase access to females by increasing the chance of meeting monokaryons. Depending on the occurrence and importance of the different sex roles in nature, and the potential tradeoffs between sex roles, selection will have shaped traits to optimize the balance of fitness in both sex roles. Our experimental evolution approach gives us the opportunity to study the effect of only the male role, as the female role was evolving minimally [76]. Selection on the monokaryotic growth and nucleus acceptance was thus relaxed, which gives the possibility to test if traits beneficial for fertilization, had deleterious pleiotropic consequences for female fitness. All strains with improved male fitness were still able to perform mating in the female role, viz. to incorporate nuclei in the mycelium. For monokaryotic growth, only one line (9P) showed reduced growth, but in other cases, no pleiotropic negative effect on female fitness traits could be shown – line 2P even showed increased growth. The apparent absence of observed trade-offs between male and female roles might be a consequence of the low cost of mating in the male role. If donating nuclei is relatively cost-free, there may be no antagonism between the two sex roles.

It remains unclear if reduced growth as observed in line 9P is caused by antagonistic pleiotropy (i.e. a male beneficial mutation with a pleiotropic deleterious effect on female fitness) or mutation accumulation of a female-deleterious allele that is neutral for mating in the male role. A male-neutral trait only present in the fertilizing nucleus is likely to be lost over time due to recombination with the non-evolving female genome. If the trait is stable over longer periods, it should either be linked to a beneficial locus, to the mating type which is always selected (linkage to mating type was not observed), or the trait itself should be beneficial in the male role. If the trait is beneficial in the male role but reduces viability in the female role (e.g. by reducing monokaryotic growth), this trait might be maintained in nature. Fertilization by spores may be a common fertilisation mechanism in nature [18,48], which skews the operational sex ratio towards males. Male beneficial traits might thus be selected even when harmful to the monokaryon, especially as the female resident nucleus in a monokaryon can compensate for a recessive allele in the fertilizing nucleus during dikaryotic vegetative growth [77,78].

Next to sexual selection of traits that improve fertilization success, natural selection can also increase fitness after the dikaryon has been established, when no clear sex roles can be distinguished any more. For instance, increased dikaryon growth might increase the number of mushrooms produced and thereby the proportion of spores. Also the functioning of the mushroom can be altered to produce more spores. Line 10P had reduced dikaryon growth rate, but this did not affect fruiting body size or spore production. Even though growth rate is often used as a proxy for fitness [79,80], it is not equal to fitness in this experiment. Mushroom formation occurs only on the early grown mycelium, and reduced growth as observed for 10P might even be advantageous, because resources used for mycelial growth can be invested in mushroom development and spore production. Except for the mushroom morphology of 6N (Fig 5), no clear changes were observed in the dikaryon phase.

## Conclusion

Increased mating success was due to a combination of increased spore formation and efficiency of establishment in the receiving mycelium. In most cases, no trade-offs were found with other fitness components.

## Acknowledgements

We acknowledge James Anderson, Arjan de Visser and Bas Zwaan for useful comments on the setup of the experiment. Bertha Koopmanschap is acknowledged for help with the qPCR assays, Luis Lugones and Karin Scholtmeijer for constructing and donating the resistant H4-8 strains, and for technical advice. We thank six anonymous reviewers for their useful comments and suggestions on previous versions of this manuscript. BPSN and DKA were financed by the Netherlands Science Organisation (ALW-NWO open competition 816.01.017; VICI; NWO 86514007).

## Data availability

Data are available at figshare doi:10.6084/m9.figshare.2065284

---

**Figure S1.**
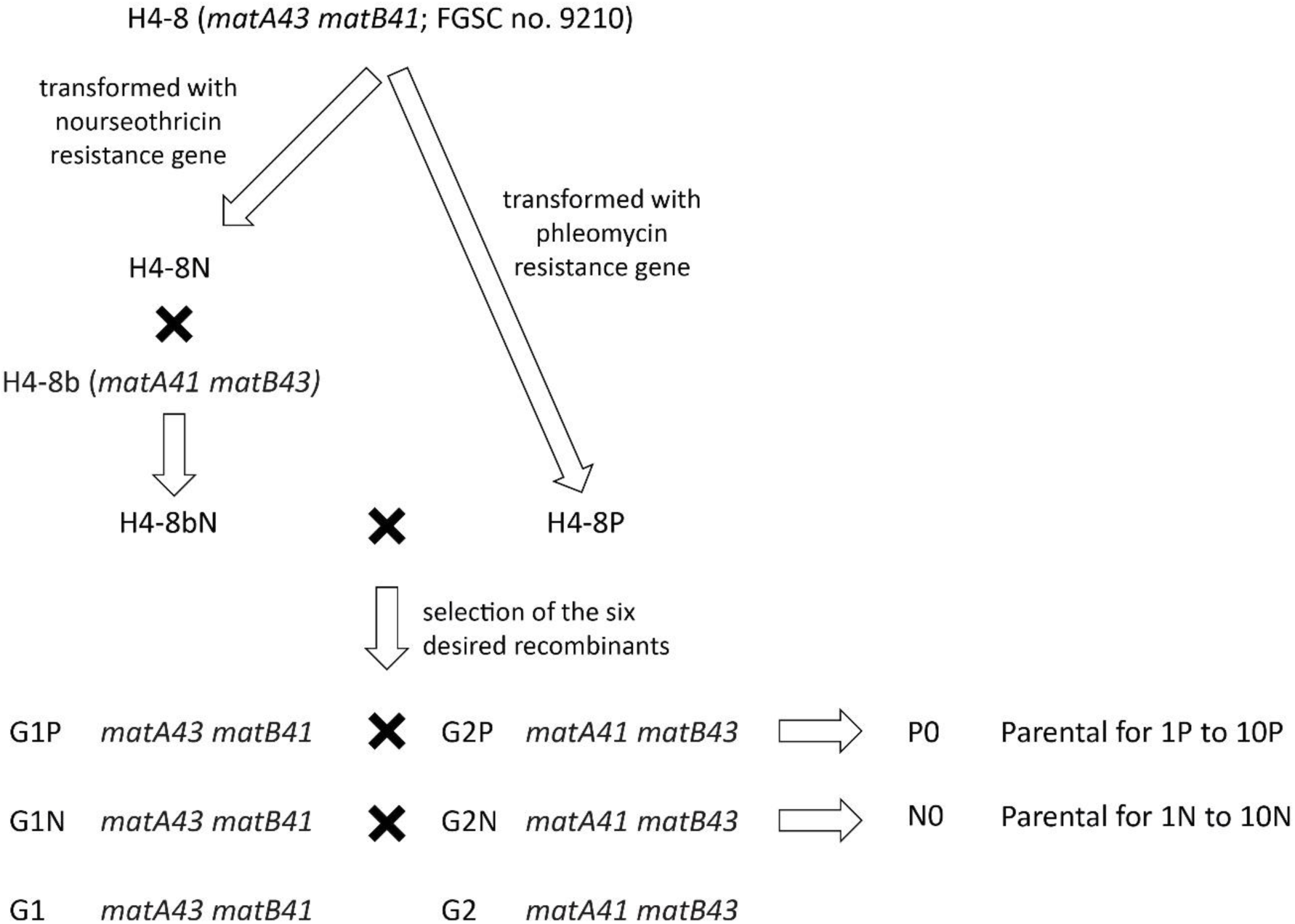
Overview of the construction of the strains used in this experiment.

**Figure S2.**
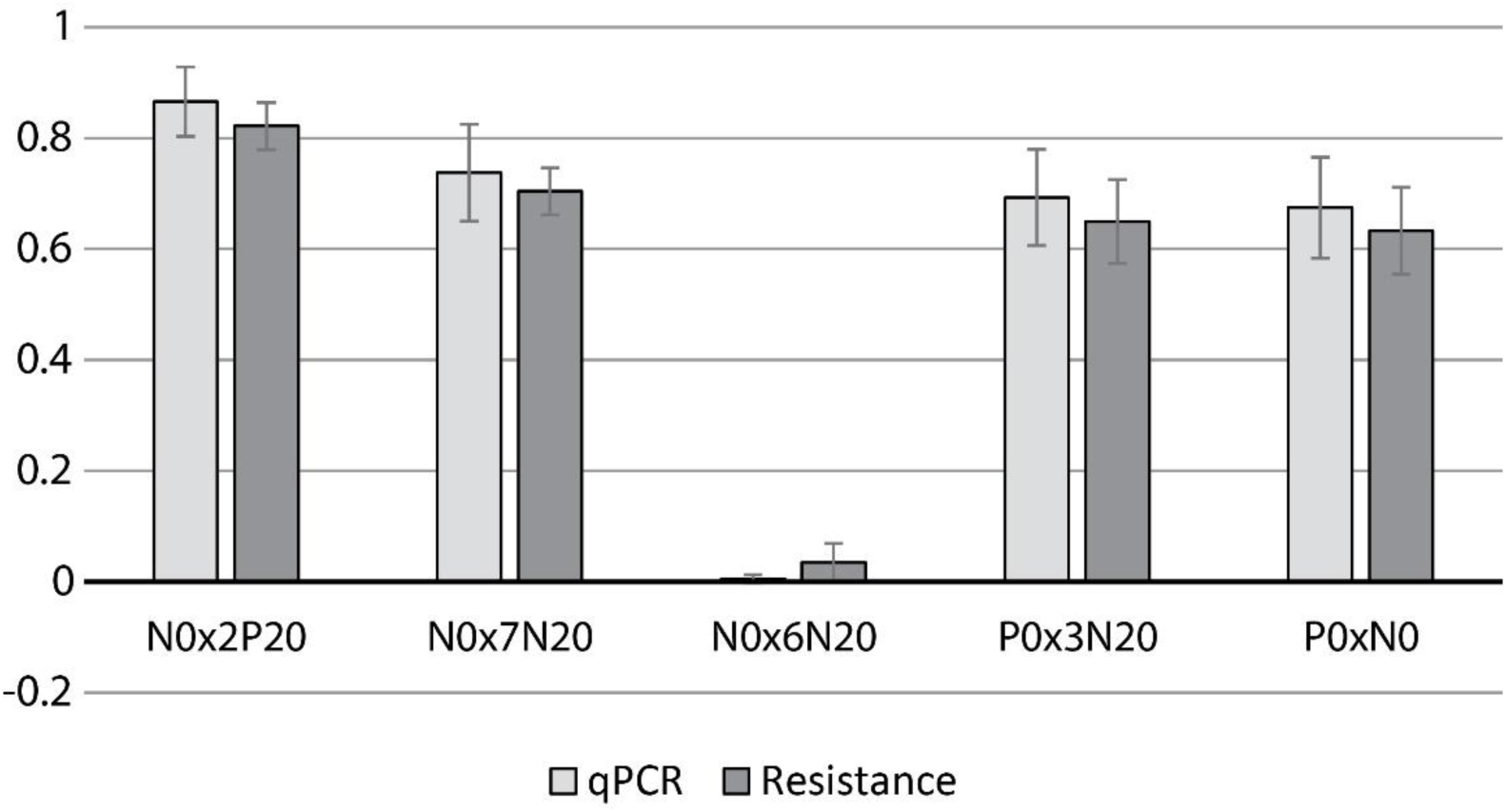
Comparison of qPCR and colony counts. The plot shows the frequency of phleomycin resistance as a proportion of all resistant strains after competition for identical samples after competition, measured from colony counts for resistance markers on plates containing triton (dark bars) and the qPCR method (light bars). See ‘Materials and Methods’ in the main text for detailed description of either methods. Mean of four replicates each. Error bars indicate standard deviation.

**Figure S3.**
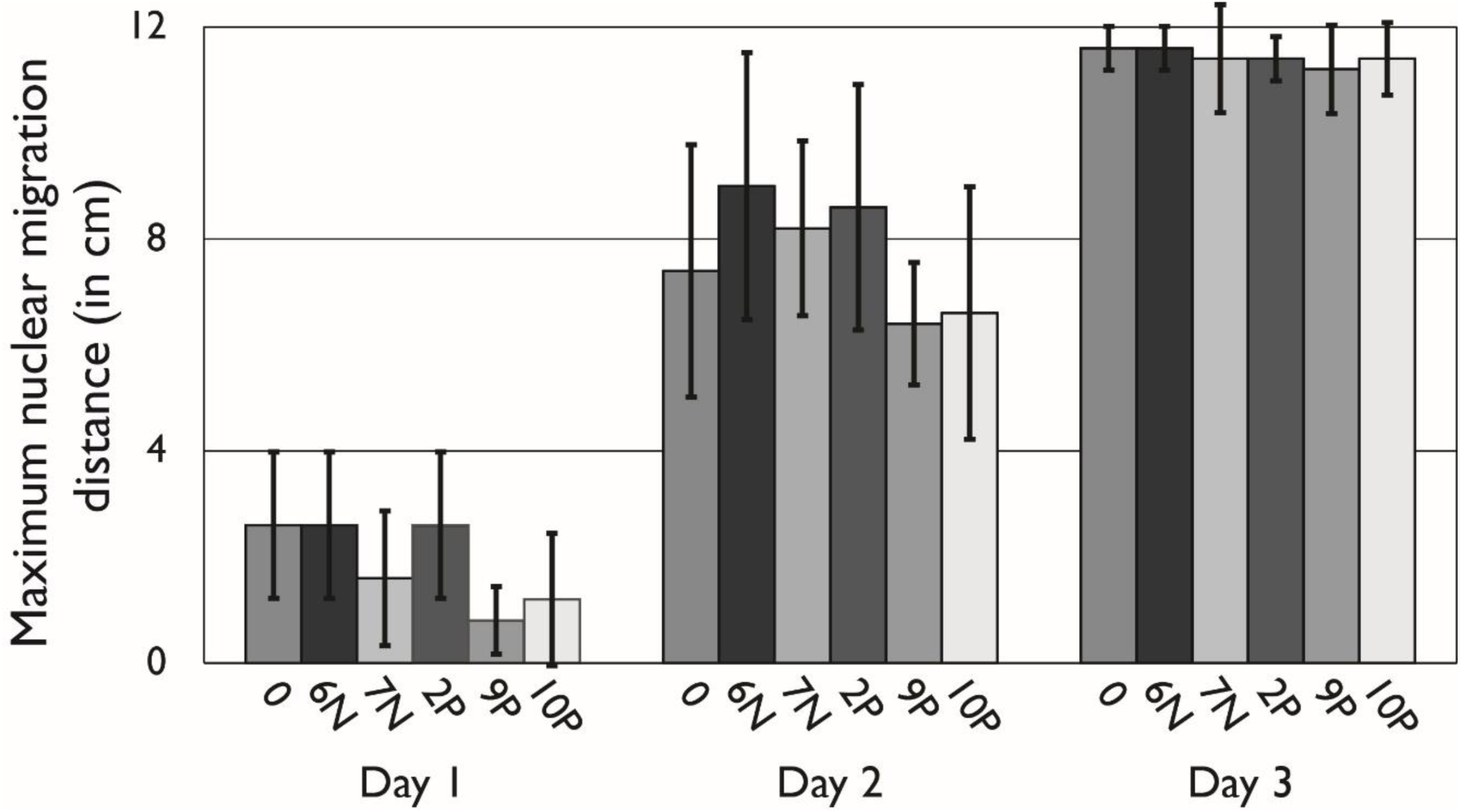
Average maximum migration distance of nuclei. Migration through a female mycelium after 1, 2 and 3 days for the parental (0) and the five evolved strains that showed changed competitive fitness at transfer 20. Measurements indicate the furthest 1 cm piece of a 12 cm ‘race track’ that was dikaryotized. Average from 5 race tracks, error bars indicate standard error.

